# MKK6 deficiency promotes cardiac dysfunction through MKK3-p38γ/δ-mTOR hyperactivation

**DOI:** 10.1101/2021.11.15.468612

**Authors:** Rafael Romero-Becerra, Alfonso Mora, Elisa Manieri, Laura Sanz, Ivana Nikolic, Ayelén M. Santamans, Valle Montalvo-Romeral, Francisco Miguel Cruz Uréndez, Maria Elena Rodríguez, Luis Leiva-Vega, Víctor Bondía, David Filgueiras-Rama, Luis Jesús Jiménez-Borreguero, José Jalife, Bárbara González-Terán, Guadalupe Sabio

## Abstract

Stress-activated p38 kinases control a plethora of functions and their dysregulation has been linked to development of steatosis, obesity, immune disorders and cancer. Therefore, they have been identified as potential targets for novel therapeutic strategies. There are four p38 family members (p38α, p38β, p38γ, and p38δ) that are activated by MKK3 and MKK6. Here we demonstrate that lack of MKK6 reduces the life span in mice. Longitudinal study of cardiac function in *Mkk6^-/-^* mice showed that young mice have cardiac hypertrophy which progresses to cardiac dilatation and fibrosis with age. Mechanistically, lack of MKK6 blunts p38α activation while causing MKK3-p38γ/δ hyperphosphorylation and increased mTOR signaling, resulting in cardiac hypertrophy. Cardiac hypertrophy in *Mkk6^-/-^* mice is reverted by knocking out either p38γ or p38δ, or by inhibiting mTOR pathway with rapamycin. In conclusion, we have identified a key role for the MKK3/6-p38γ/δ pathway in the development of cardiac hypertrophy, which has important implications for the clinical use of p38α inhibitors in the long-term treatment since they might result in cardiotoxicity.

**eLife’s Review Process:** eLife works to improve the process of peer review so that it more effectively conveys the assessment of expert reviewers to authors, readers and other interested parties. In the future we envision a system in which research is first published as a preprint and the outputs of peer review are the primary way research is assessed, rather than journal title.

Our editorial process produces two outputs: i) an assessment by peers designed to be posted alongside a preprint for the benefit of the readers; ii) detailed feedback on the manuscript for the authors, including requests for revisions and suggestions for improvement.

Therefore we want to change how we construct and write peer reviews to make them useful to both authors and readers in a way that better reflects the work you put into reading and thinking about a paper.

eLife reviews now have three parts:

- An **evaluation summary** (in two or three sentences) that captures the major conclusions of the review in a concise manner, accessible to a wide audience.
- A **public review** that details the strengths and weaknesses of the manuscript before you, and discusses whether the authors’ claims and conclusions are justified by their data.
- A set of private **recommendations for the authors** that outline how you think the science and its presentation could be strengthened.

All three sections will be used as the basis for an eLife publishing decision, which will, as always, be made after a consultation among the reviewers and editor. Each of the **public reviews** will be published (anonymously) alongside the preprint, together with a response from the authors if they choose. In the case of papers we reject after review, the authors can choose to delay posting until their paper has been published elsewhere.

If this is your first time going through this new process, we ask that you take some time to read our [Reviewer Guide](https://submit.elifesciences.org/cgi-bin/main.plex?form_type=display_rev_instructions#writing), which discusses how we see each section will be used, what it should contain, and what we hope it accomplishes. And we remind you that, with the shift of reviews from private correspondence to public discourse, it is more important than ever that reviews are written in a **<u>clear and constructive manner</u>** appropriate for a public audience and mindful of the impact language choices might have on the authors.

## INTRODUCTION

Cardiac hypertrophy is an adaptive response of the heart to hemodynamic stress that can be physiologic (e.g., pregnancy or exercise) or pathological (e.g., hypertension or valvular disease). Physiological cardiac hypertrophy is accompanied with a normal or even enhanced cardiac function, while pathological forms of hypertrophy are accompanied by myocardial dysfunction and fibrosis and represent a risk factor for ventricular arrhythmias and sudden cardiac death (Maillet *et al*, 2013; Nakamura & Sadoshima, 2018; Oldfield *et al*, 2020).

Initially, cardiac hypertrophy is induced as a compensatory response to preserve cardiac function under stressful conditions, a process known as adaptive cardiac hypertrophy. However, if the pathological stimulus is maintained, this adaptive cardiac hypertrophy will eventually lead to the development of pathological cardiac hypertrophy and heart failure (Nakamura & Sadoshima, 2018; Oldfield *et al*., 2020). The form of cardiac hypertrophy developed will depend on the type of the hypertrophic stimuli, the duration of the stimuli and the downstream signaling involved (Nakamura & Sadoshima, 2018; Oldfield *et al*., 2020; Shimizu & Minamino, 2016). Several signalling pathways known to promote physiological cardiac hypertrophic growth, when persistently activated have been found to drive pathological hypertrophy and cardiac dysfunction (Heineke & Molkentin, 2006; Maillet *et al*., 2013; Nakamura & Sadoshima, 2018; Porrello *et al*, 2008). For instance, IGF1 or Akt transgenic mice develop proportionately enlarged hearts with initially normal cardiac function, which over time progress to pathological hypertrophy with impaired cardiac function (Delaughter *et al*, 1999; Shiojima *et al*, 2005).

Stress-inducing stimuli in the heart activate several mitogen-activated protein kinases (MAPKs) including the p38 family. p38 kinases control a wide range of processes and their dysregulation has been linked to numerous diseases, making them a promising pharmacological target for therapeutic use (Canovas & Nebreda, 2021). This family consists of four isoforms: α, β, γ, and δ, with p38α having been the most broadly studied, whereas knowledge of the other p38 isoforms has been limited by a reduced availability of isoform-specific reagents. Our previous work showed that p38γ and p38δ are expressed in the heart and participate in the cardiac hypertrophic response. We have shown that p38γ/δ mediate early postnatal cardiac hypertrophy by promoting mTOR-induced cell growth (Gonzalez-Teran *et al*, 2016).

An essential feature of both physiological and pathological hypertrophy is increased protein synthesis, critically regulated by the mammalian target of rapamycin (mTOR) pathway mainly through the phosphorylation of its downstream substrates. Activation of mTOR signaling is increased during postnatal cardiac development (Gonzalez-Teran *et al*., 2016) as well as in the hearts of transgenic mouse models suffering from physiological cardiac hypertrophy (McMullen *et al*, 2004a; McMullen *et al*, 2004b; Shioi *et al*, 2000; Shioi *et al*, 2003). Moreover, the specific mTOR inhibitor rapamycin attenuates and reverses cardiac-overload-induced pathological hypertrophy (Shioi *et al*., 2003). Conversely, mTOR pathway activation mediated by p38γ and p38δ MAPKs has been implicated in the control of postnatal cardiac hypertrophic growth and angiotensin-II-induced cardiac hypertrophy (Gonzalez-Teran *et al*., 2016).

Here, we demonstrate that in the heart MKK3 activates p38γ/δ, whereas MKK6 activates p38α. Furthermore, we find that *Mkk6^-/-^* mice exhibit cardiac hypertrophy caused by hyperactivation of the MKK3-p38γ/δ axis, which progresses to a pathological cardiac hypertrophy phenotype with age. Our results have important implications for the clinical use of p38α inhibitors in the long-term treatment since they might result in cardiotoxicity.

## RESULTS

### MKK6-deficient mice die prematurely

Several studies have addressed the role of p38 signaling in homeostasis and disease (Nikolic *et al*, 2020; Romero-Becerra *et al*, 2020). While mice lacking both MKK3 and MKK6 die in mid-gestation with mutant embryos demonstrating abnormalities of the placenta and embryonic vasculature (Brancho *et al*, 2003), mice individually lacking MKK3 or MKK6 are viable and fertile, suggesting partial functional redundancy (Lu *et al*, 1999; Tanaka *et al*, 2002; Wysk *et al*, 1999). However, the role of the p38 pathway in aging remains incompletely understood. Therefore, we examined mice harboring germline deletion of *Mkk6* (*Mkk6^-/-^*) (Tanaka *et al*., 2002) at advanced age. Mice lacking *Mkk6* have reduced body weight compared to age-matched wild type (WT) animals (Figure 1A, B), which can be partially explained by a dramatic reduction in white adipose tissue (Figure 1C). This agrees with previous studies demonstrating that *Mkk6^-/-^* mice are protected against diet-induced obesity with increase browning of the epididymal white adipose tissue (eWAT) (Matesanz *et al*, 2017).

**Figure 1.**
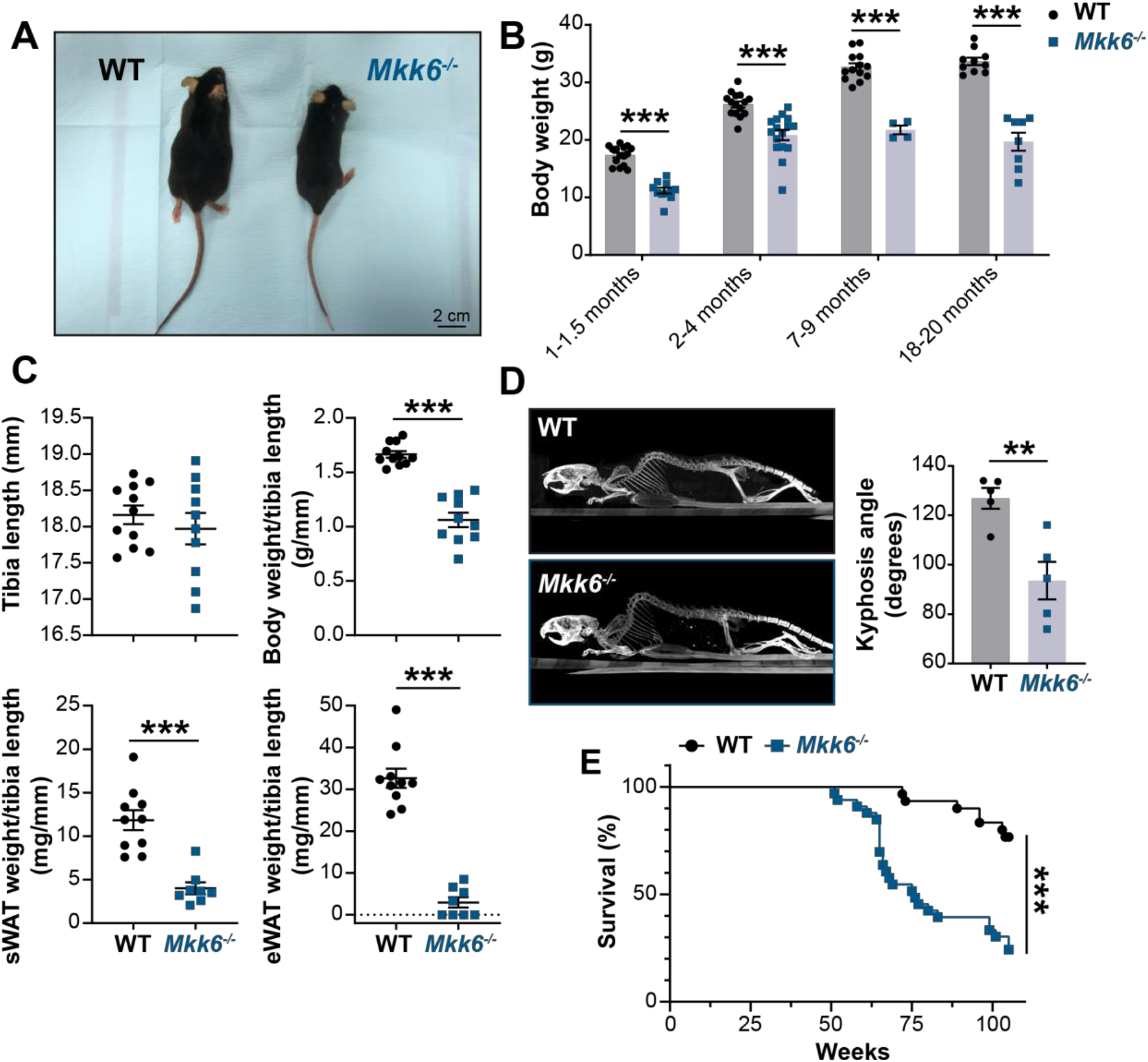
*Mkk6^-/-^* mice show a reduced survival age. **(A)** Representative picture of 19-month-old WT and *Mkk6^-/-^* male mice. Scale bar: 2 cm. **(B)** Body weight of WT (*n*=10-15) and *Mkk6^-/-^* (*n*=4-16) mice over the indicated age period. 2-way ANOVA followed by Sidak’s post test. (**C**) Tibia length and body weight, subcutaneous white adipose tissue (sWAT) and epidydimal white adipose tissue (eWAT) to tibia length ratio from 20-month-old WT (*n*=10-11) and *Mkk6^-/-^* (*n*=8-10) mice. Unpaired *t*-test or Mann-Whitney test. (**D**) Representative CT scan images and quantification of the column kyphosis angle of 19-week-old WT (*n*=5) and *Mkk6^-/-^* (*n*=5) mice. Unpaired *t*-test. (**E**) Kaplan-Meier survival plot of age-related mortality in WT (*n*=30) and *Mkk6^-/-^* (*n*=33) mice. An endpoint of 105 weeks was chosen to avoid a severe worsening of the mice health. Gehan-Breslow-Wilcoxon test. Data in B-D are mean ± SEM. \*\**P*<0.01; \*\*\**P*<0.001.

Additionally, these mice exhibit an abnormal posture characterized by a hunched position and the development of thoracic kyphosis and severe ataxia (Figure 1D and Figure 1 – video supplement 1). As a consequence of all these phenotypic alterations, *Mkk6^-/-^* mice suffer premature death, the first mice dying at 51 weeks of age with a median life span of 76 weeks (Figure 1E).

### MKK6-deficient mice develop increased age-related cardiac dysfunction

The downstream kinases of MKK6 have been implicated in major cardiovascular abnormalities during development. Combined deletion of p38α and p38β results in cardiac defects during embryonic development (del Barco Barrantes *et al*, 2011), whereas p38γ/δ deficient mice exhibit reduced cardiomyocyte hypertrophic growth and smaller hearts (Gonzalez-Teran *et al*., 2016). This prompted us to speculate that cardiac abnormalities could be one of the underlying causes of premature death of *Mkk6^-/-^* mice. Echocardiographic analyses of 12 to 14-month-old mice demonstrated eccentric hypertrophy in *Mkk6^-/-^* mice compared to control mice, as detected by thinning of the left ventricle (LV) wall, as well as increased left ventricular internal diameter (LVID) and left ventricular volume, especially during the systole (Figure 2A). Cardiac enlargement compromised systolic function, evidenced by a decreased in the ejection fraction and fractional shortening. However, the diastolic function appeared to be maintained, with a normal E/A wave velocities ratio and isovolumetric relaxation time (Figure 2B). Moreover, *Mkk6^-/-^* mice exhibit bradycardia (Figure 2C). We performed picrosirius red staining for collagen and quantified positive areas in serial histologic sections from *Mkk6^-/-^* and WT hearts and found cardiac fibrotic lesions in *Mkk6^-/-^* old mice (Figure 2D). To discard hypertension as a possible contributor of the cardiac dysfunction we evaluated blood pressure in these animals. *Mkk6^-/-^* mice did not present differences in blood pressure compared to age-matched controls (Figure 2E).

**Figure 2.**
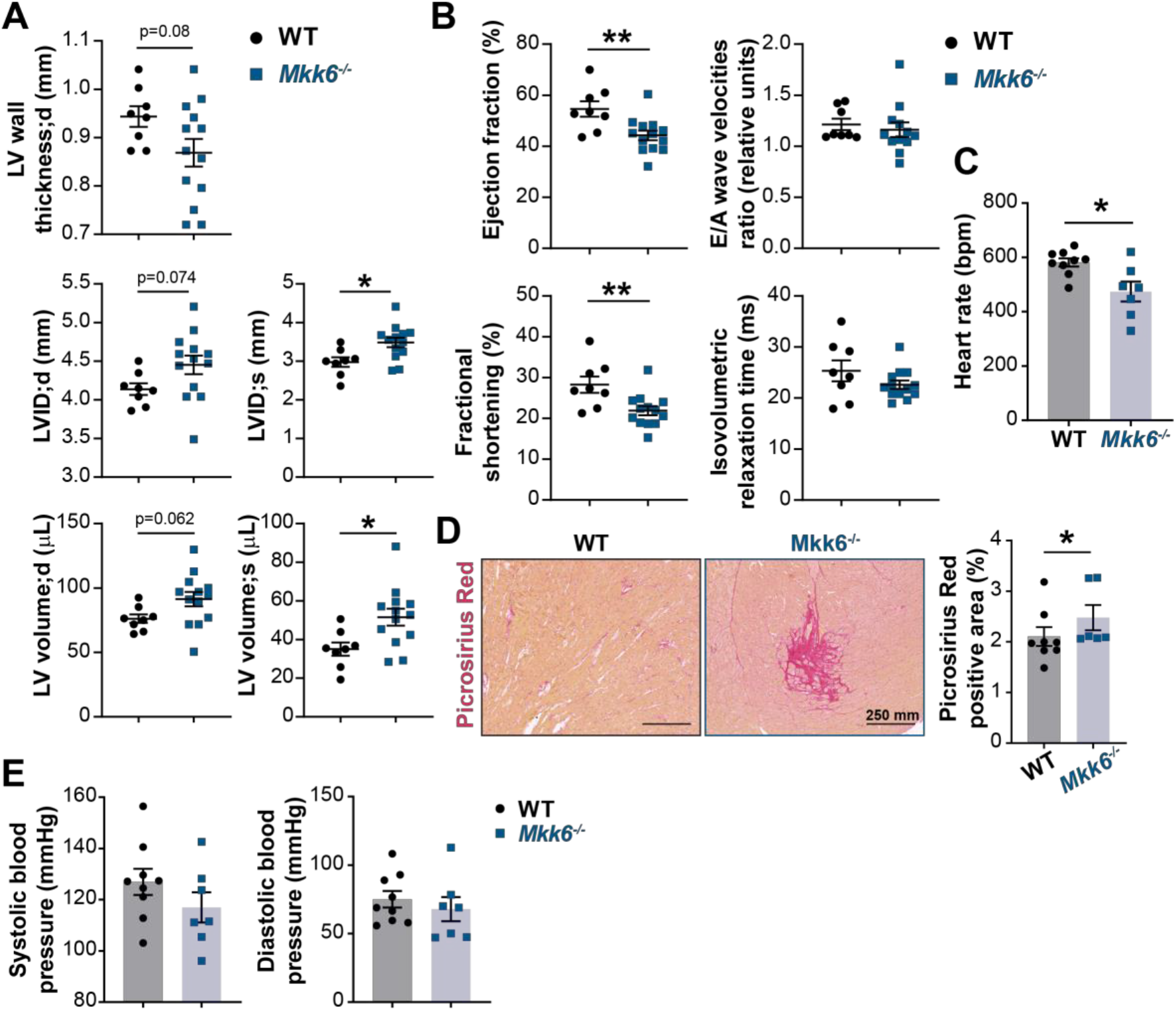
MKK6 deficiency promotes cardiac dysfunction at advanced ages. (**A**, **B**) Echocardiography parameters related to left ventricle (LV) dimensions (A) and contractility (B) in 12 to 14-month-old WT (*n*=8) and *Mkk6^-/-^* (*n*=13). Each dot corresponds to an individual animal. LV wall thickness; d (left ventricle wall thickness in diastole), LVID;d (left ventricular internal diameter in diastole), LVID;s (left ventricular internal diameter in systole), LV volume;d (left ventricular volume in diastole), LV volume;s (left ventricular volume in systole). Unpaired *t*-test. (**C**) Heart rate in conscious 18-month-old WT (*n*=9) and *Mkk6^-/-^* (*n*=7) mice. bpm (beats per minute). Unpaired *t*-test. (**D**) Picrosirius red staining and quantification of cardiac fibrosis in 23 to 24-month-old WT (*n*=8) and *Mkk6^-/-^* (*n*=6) mice. Mann-Whitney test. Scale bars: 250 μm. (**E**) Systolic and diastolic blood pressure in measured in conscious 18-month-old WT (*n*=9) and *Mkk6^-/-^* (*n*=7) mice. Unpaired *t*-test. Data in A-E are mean ± SEM. \**P*<0.05; \*\**P*<0.01.

### Young MKK6-deficient mice present cardiac hypertrophy

Cardiac dysfunction may result from an initial compensated cardiac hypertrophy that with time becomes pathological (Nakamura & Sadoshima, 2018). Echocardiographic analysis at 9 weeks of age demonstrated cardiac hypertrophy in MKK6-deficient animals when compared with controls, as detected by measures of left ventricular mass, interventricular septal thickness, left ventricular posterior wall thickness, and left ventricular internal diameter (Figure 3A). However, cardiac enlargement did not compromise systolic function or diastolic function (given by the ejection fraction and the E/A wave velocity ratio, respectively) but was accompanied by increases in stroke volume and cardiac output (Figure 3B). Gross anatomic and histologic analyses confirmed these non-invasive findings as MKK6-deficient hearts were larger than WT controls when normalized to tibia length (TL) (Figure 3C&D), a difference that was not apparent at 4 weeks of age. Serial analysis of heart weight (HW)/TL over 15 weeks demonstrated enhanced cardiac growth in *Mkk6^-/-^* mice (Figure 3D). The progressive increase in size of *Mkk6^-/-^* hearts correlated with increased cardiomyocyte cross-sectional area, consistent with enhanced hypertrophic growth (Figure 3E). Hypertension was excluded as a possible contributor to this increased growth as *Mkk6^-/-^* mice demonstrate reduced systolic blood pressures when compared to age-matched controls (Figure 3 – figure supplement 1A). Fibrosis and reactivation of a “fetal gene program” (e.g., *Nppa, Nppb, Acta2, Myh7*) are hallmark features of pathologic hypertrophy (Bernardo *et al*, 2010). Histological cardiac examination revealed no evidence of fibrosis in *Mkk6^-/-^* heart sections (Figure 3 – figure supplement 1B). We also found no difference in the expression of fibrotic genes including *Colla1, Col3a1, and Fn* or markers of the fetal gene program (Figure 3 - figure supplement 1C, D). No meaningful changes were visible for *Nppa* or *Nppb* (Figure 3 – figure supplement 1D). These observations suggest that at 9 weeks of age *Mkk6*^-/-^ hearts show a non-pathological cardiac hypertrophy.

**Figure 3.**
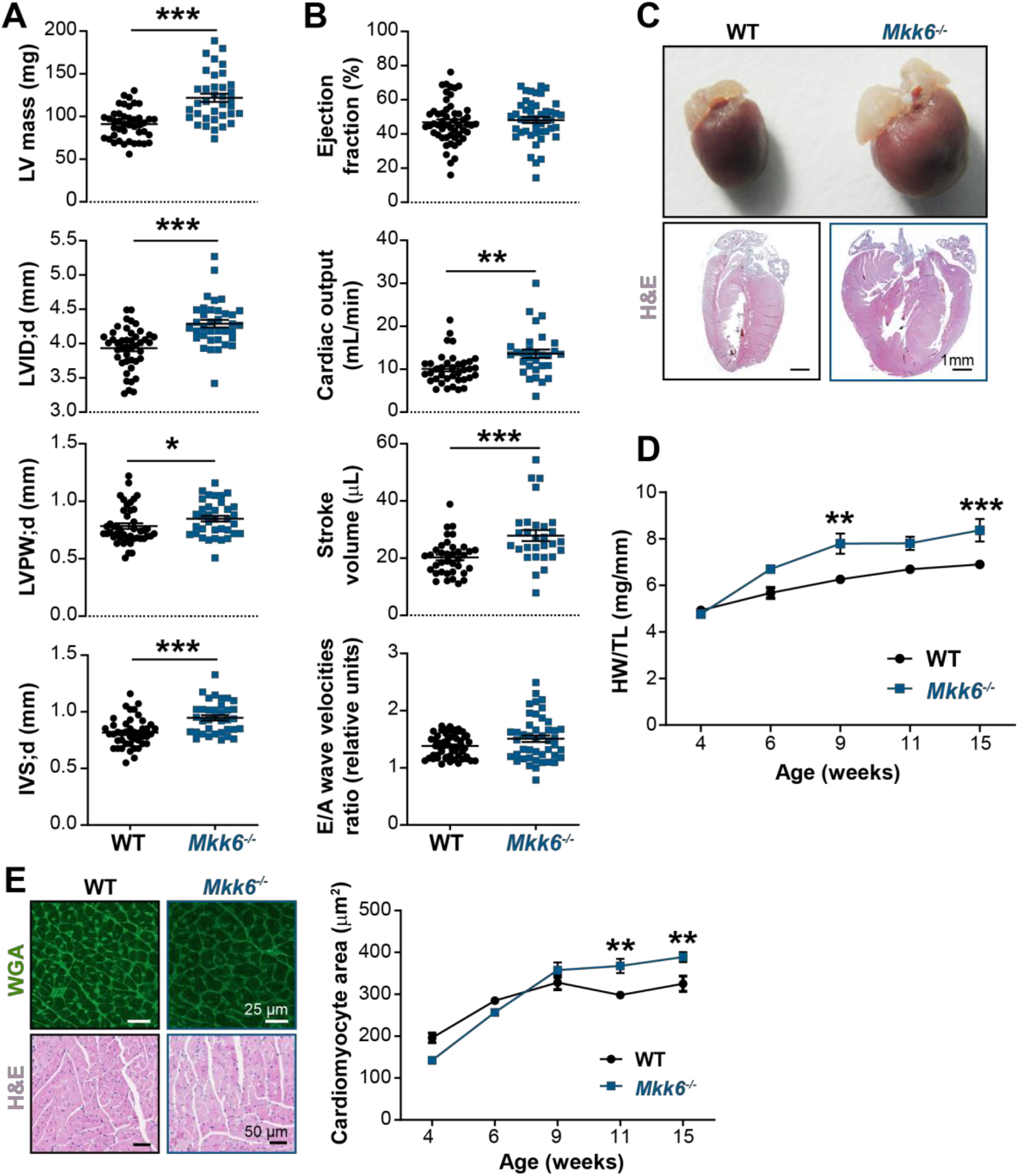
Young MKK6-deficient hearts are hypertrophic with preserved cardiac function. (**A, B**) Echocardiography parameters related to left ventricle (LV) dimensions (A) and contractility (B) in 9-week-old WT (*n*=37-54) and *Mkk6^-/-^* (*n*=29-46) mice. Each dot corresponds to an individual mouse. Mean ± SEM are shown as well. LV Mass (left ventricular mass), LVID;d (left ventricular internal diameter in diastole), LVPW;d (left ventricular posterior wall in diastole), IVS;d (inter-ventricular septum in diastole). Unpaired *t*-test or Mann-Whitney test. **(C**) Representative whole hearts and cardiac longitudinal sections stained with hematoxylin & eosin (H&E) from 9-week-old WT and *Mkk6^-/-^* mice. Scale bars: 1 mm. (**D**) Heart weight to tibia length ratio (HW/TL) of WT (*n*=4-15) and *Mkk6^-/-^* (*n*=5-14) over the indicated age period. 2-way ANOVA followed by Sidak’s post test. (**E**) *Top:* Representative FITC wheat germ agglutinin (FITC-WGA)-stained heart sections from 9-week-old WT and *Mkk6^-/-^* mice and quantification of cardiomyocyte cross-sectional area over time (right graph, WT *n*=4-5; *Mkk6^-/-^ n*=4-6, 2-way ANOVA followed by Sidak’s post test). Scale bars: 25 μm. *Bottom*. Representative H&E-stained heart sections. Scale bars: 50 μm. Data in A, B, D and E are mean ± SEM. \**P*<0.05; \*\**P*<0.01; \*\*\**P*<0.001.

To confirm the MKK6 autonomous effect in cardiomyocytes we employed a murine conditional MKK6 allele (*Mkk6^LoxP^)^23^* and two cardiomyocyte Cre-expressing lines (*MCK-Cre* (Bruning *et al*, 1998) and *αMHC-Cre* (McFadden *et al*, 2005)) to assess the consequences of MKK6 genetic ablation in postnatal cardiomyocytes. MCK-Cre is active in striated muscle, and the *MCK-Cre*; *Mkk6^LoxP/LoxP^* (Mkk6^MCK–KO^) mice demonstrated specific and efficient deletion of MKK6 in the heart and skeletal muscle tissues, but not in spleen or liver (Figure 4 – figure supplement 1A, B). Importantly, the cardiac phenotype of 9-weeks-old Mkk6^MCK–KO^ mice resembled that of *Mkk6^-/-^* animals (Figure 4A-C). Similar results were obtained with *αMHC-Cre, Mkk6^LoxP/LoxP^* (Mkk6^αMHC–KO^) mice (Figure 4D-F). These data collectively confirm that cardiomyocyte MKK6 controls heart growth.

**Figure 4.**
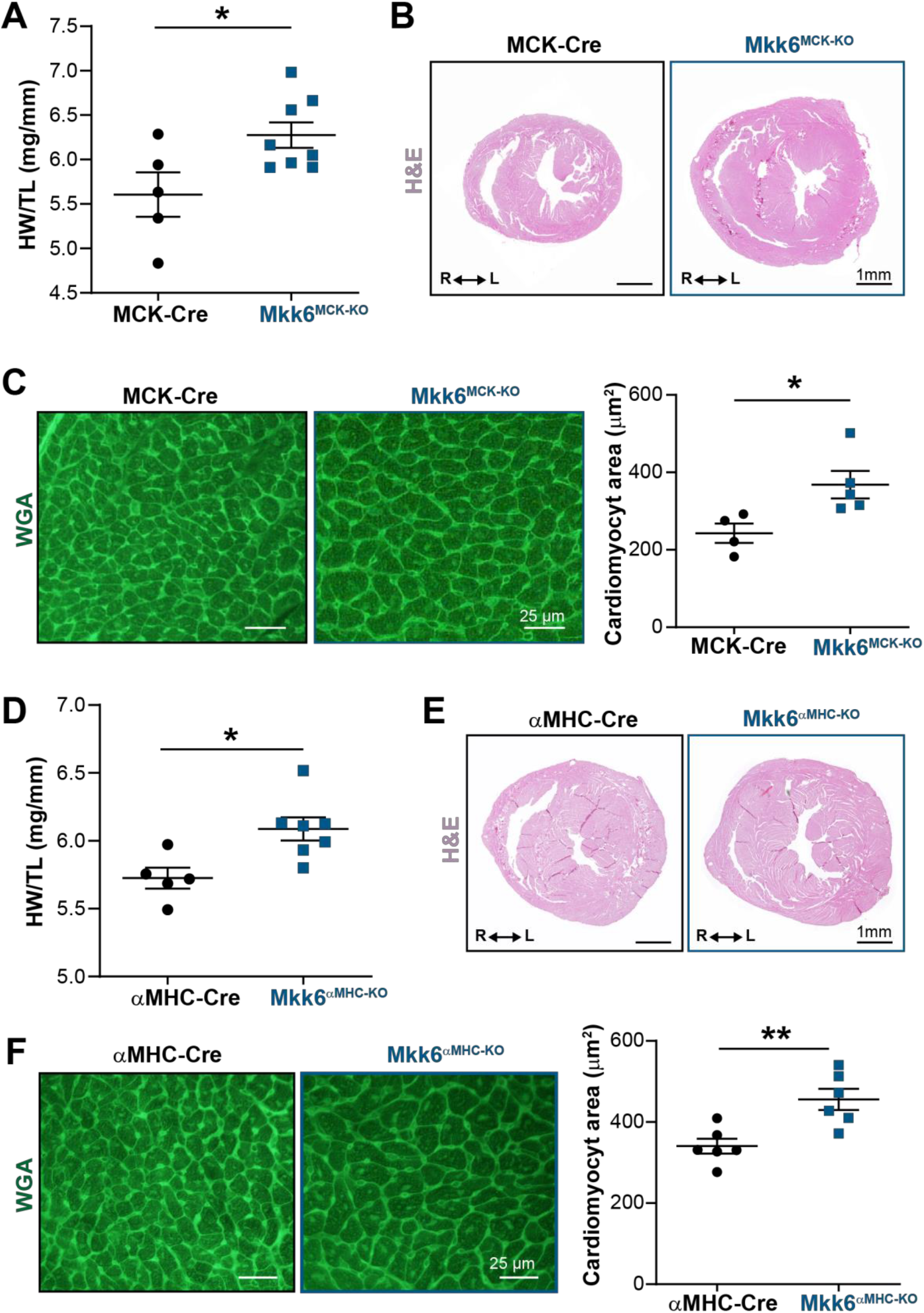
Cardiac MKK6 controls postnatal heart growth. (**A**) Heart weight to tibia length ratio (HW/TL) in 9-week-old MCK-Cre (*n*=5) and Mkk6^MCK-KO^ (*n*=8) mice. Unpaired *t*-test. (**B**) H&E-stained transverse cardiac sections from control (MCK-Cre) and Mkk6^MCK-KO^ mice. Scale bars: 1 mm. (**C**) Representative FITC wheat germ agglutinin (FITC-WGA) staining and corresponding quantification of cardiomyocyte cross-sectional area in MCK-Cre (*n*=4) and Mkk6^MCK-KO^ (*n*=5) mice. Unpaired *t*-test. Scale bars: 25 μm. (**D**) Heart weight to tibia length ratio in 9-week-old αMHC-Cre (*n*=5) and MKK6^αMHC-KO^ (*n*=7) mice. Unpaired *t*-test. (**E**) H&E-stained transverse heart sections from αMHC-Cre and MKK6^αMHC-KO^ mice. Scale bars: 1 mm. (**F**) Representative FITC-WGA staining and corresponding quantification of cardiomyocyte cross-sectional area in αMHC-Cre (*n*=6) and MKK6^αMHC-KO^ (*n* 6) mice. Unpaired *t*-test. Scale bars: 25 μm. Data in A, C, D and F are mean ± SEM. \**P*<0.05; \*\**P*<0.01.

### MKK6-deficient hearts have increased MKK3-p38γ/δ activation

MKK6 is a critical upstream activator of p38 MAPKs, but its specificity for individual p38 family isoforms is not well established. We assessed the relative levels of phosphorylated p38 isoforms by immunoprecipitation in *Mkk6^-/-^* hearts, which demonstrated hyperphosphorylation of p38γ and p38δ (Figure 5A) with a simultaneous reduction in phosphorylation of p38α (Figure 5B). Immunoblot analysis also revealed increased levels of phosphorylated MKK3, the other main p38 upstream activator, in *Mkk6^-/-^* hearts (Figure 5C). This observation suggested that increased phosphorylation of p38γ and p38δ in the *Mkk6^-/-^* hearts resulted from MKK3 activation. Accordingly, phosphorylation of p38γ and p38δ was strongly reduced in MKK3-deficient mice, whereas p38α phosphorylation was not changed appreciably (Figure 5D). In addition, hearts of *Mkk3^-/-^* mice were smaller at 9 weeks of age when compared with age-matched WT controls (Figure 5E), a finding consistent with the previously described roles of p38γ and p38δ in promoting postnatal cardiac hypertrophic growth(Gonzalez-Teran *et al*., 2016). Taken together, these observations indicate that MKK6 primarily targets p38α and MKK3 p38γ and p38δ. In addition, they suggest that the MKK6 deficiency leads to cardiac hypertrophy via activation of MKK3 and p38γ and p38δ.

**Figure 5.**
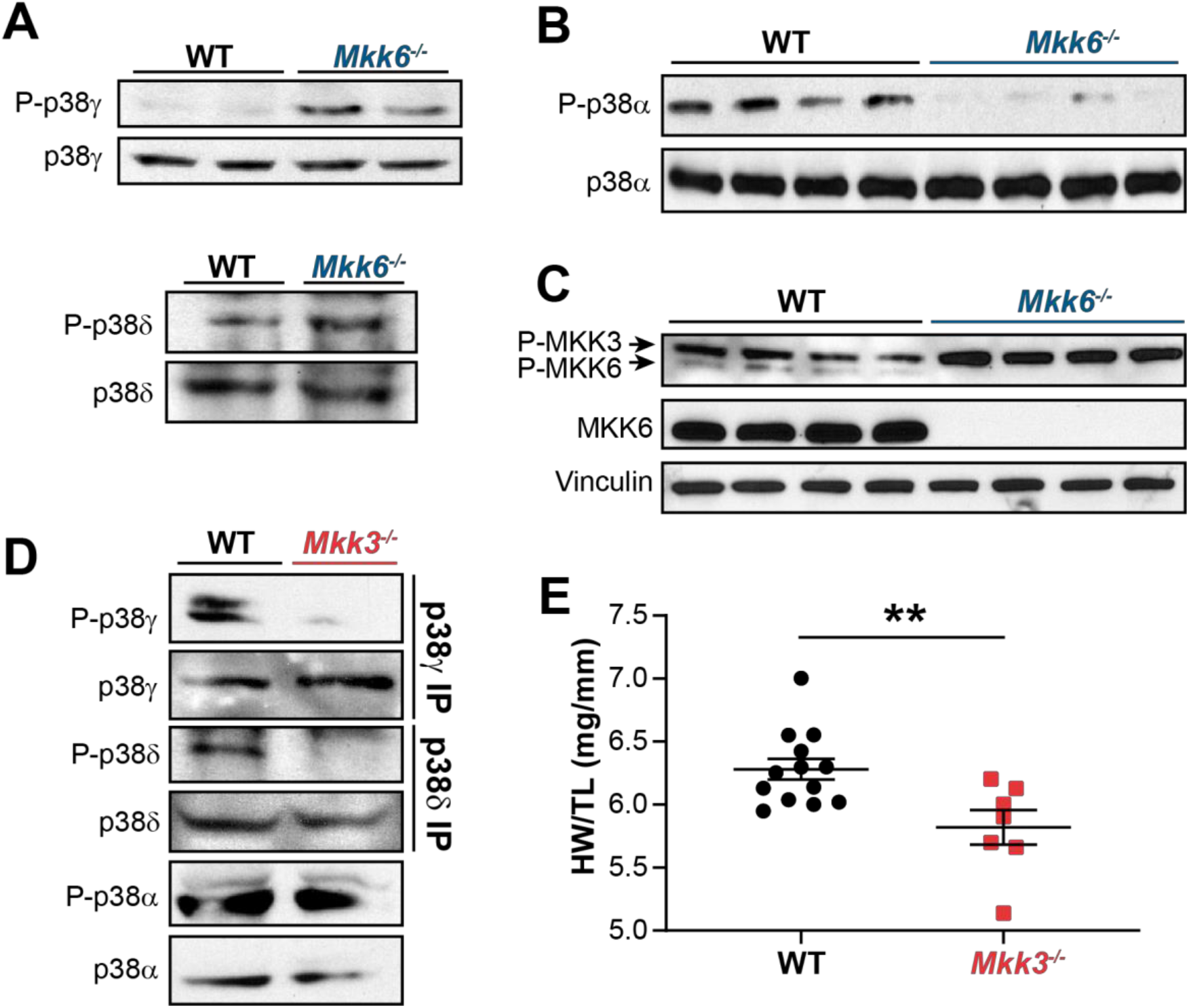
MKK6 is necessary for p38α phosphorylation and MKK3 for p38γ and p38δ phosphorylation in the heart. (**A)** Western blot analysis of the phosphorylation and amount of p38γ and δ immunoprecipitated from heart lysates from 9-week-old WT and *Mkk6^-/-^* mice. (B) Immunoblot analysis of p38α phosphorylation and protein amount in WT and *Mkk6^-/-^* mice. (**C**) Phosphorylation and protein levels of MKK3 and MKK6 in heart lysates from WT and *Mkk6^-/-^* mice. (**D**) Immunoprecipitation analysis of the phosphorylation and protein amounts of p38α, p38γ and p38δ isoforms in heart lysates from 9-week-old WT and *Mkk3^-/-^* mice. (**E**) Heart weight to tibia length ratio in WT (*n*=13) and *Mkk3^-/-^* (*n*=7) mice at 9 weeks of age. Data in E are mean ± SEM. (*n*=7-13). \*\**P*<0.01 (Unpaired *t*-test).

### Hypertrophy in MKK6-deficient hearts is mediated by p38γ/δ

To confirm that enhanced hypertrophic growth in MKK6-deficient mice is mediated by modulation of p38γ/δ activation, we introduced a deletion of p38γ in the context of MKK6 deficiency. The double mutant combination (*Mkk6^-/-^*; p38γ*^-/-^*) rescued normal cardiac growth, as the heart sizes and cardiomyocyte cross-sectional areas of the double-mutants were equivalent to those of WT controls (Figure 6A-C). We further demonstrated the requirement for p38δ to be cell-autonomous in striated muscle using a p38δ conditional allele (*Mapk13^LoxP^*). MKK6-deficient hearts lacking p38δ in their myocytes (*Mkk6^-/-^*; *MCK-Cre*; *p38δ^LoxP/LoxP^* hearts) were similar in size to those of mice lacking p38δ in their myocytes (*MCK-Cre*; *p38δ ^LoxP/LoxP^* (p38δ^MCK–KO^)), with no appreciable increase in cardiomyocyte cross-sectional area (Figure 6D-F). Collectively, these data demonstrate that enhanced cardiac hypertrophic growth in MKK6-deficient mice is mediated by hyperactivation p38γ/δ of signaling in striated muscle.

**Figure 6.**
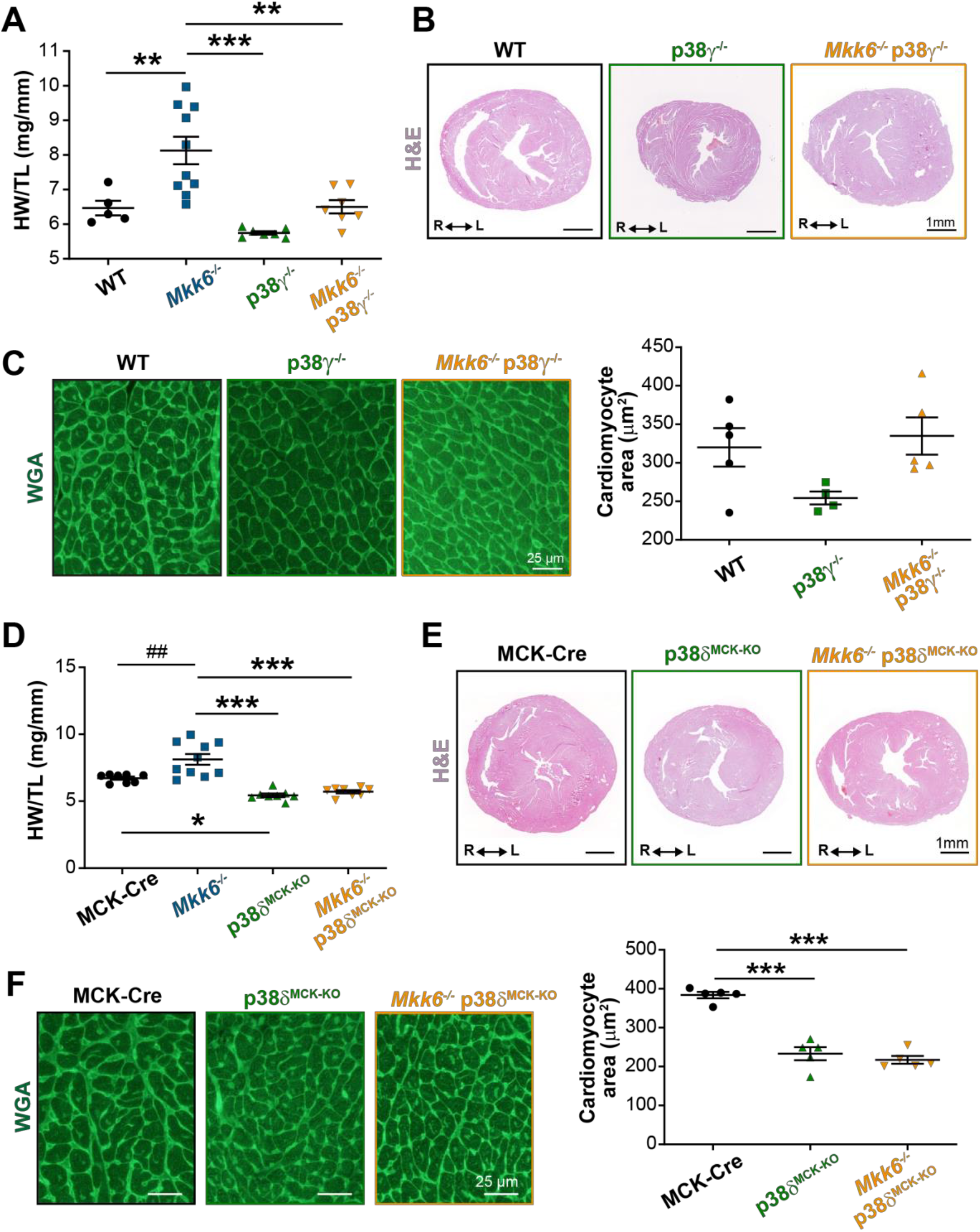
Loss of p38γ/δ in cardiomyocytes rescues the cardiac hypertrophy induced by MKK6 deficiency. All phenotypes shown come from 9-week-old mice. **(A**) Heart weight to tibia length ratio in WT (*n*=5), *Mkk6^-/-^* (*n*=10), p38γ^-/-^ (*n*=7) and *Mkk6^-/-^* p38γ^-/-^ (*n*=7) mice. 1-way ANOVA followed by Tukey’s post test. (B) Representative H&E-stained transverse heart sections from WT, p38γ^-/-^ and *Mkk6^-/-^* p38γ^-/-^ mice. Scale bars: 1 mm. (**C**) Representative FITC-WGA staining and corresponding quantification of cardiomyocyte cross-sectional area in WT (*n*=5), p38γ^-/-^ (*n*=4) and *Mkk6^-/-^* p38γ^-/-^ (*n*=5) mice. Scale bars: 25 μm. 1-way ANOVA followed by Tukey’s post test. **(D**) Heart weight to tibia length ratio in MCK-Cre (*n* 8), *Mkk6^-/-^* (*n*=10), p38δ^MCK-KO^ (*n*=8), and *Mkk6^-/-^* p38δ^MCK-KO^ (*n* 8) mice. Kruskal-Wallis test with Dunn’s post-test (##*P*<0.01 Mann-Whitney test). (**E**) Representative H&E-stained transverse heart sections from MCK-Cre, p38δ^MCK-KO^, and *Mkk6^-/-^* p38δ^MCK-KO^ mice. Scale bars: 1 mm. (**F**) Representative FITC-WGA staining and corresponding quantification of cardiomyocyte cross-sectional area in MCK-Cre (*n*=5), p38δ^MCK-KO^ (*n*=5), and *Mkk6^-/-^* p38δ^MCK-KO^ (*n*=5) mice. 1-way ANOVA followed by Tukey’s post test. Scale bars: 25 μm. The same data from *Mkk6^-/-^* mice was used in (A) and (D). Means ± SEM are shown. \**P*<0.05; \*\**P*<0.01; \*\*\**P*<0.001; ##*P*<0.01.

### mTOR pathway hyperactivation mediated hypertrophy in MKK6-deficient hearts

The p38γ and p38δ isoforms have previously been demonstrated to promote cardiac hypertrophic growth through activation of mTOR signaling (Gonzalez-Teran *et al*., 2016). As expected from that finding, immunoblot analysis of *Mkk6^-/-^* mouse hearts showed an increase in mTOR pathway activation (Figure 7A). As protein synthesis represents a key target of the mTOR signaling pathway and is critical for cardiomyocyte hypertrophic growth, we assessed eukaryotic initiation/elongation factors by immunoblot analyses. We found an overall increase in translational activation in *Mkk6^-/-^* hearts relative to WT (Figure 7B). We corroborated these findings by analyzing puromycin incorporation into newly synthesized peptides in WT and *Mkk6^-/-^* hearts, which demonstrated greater puromycin-labeling (Figure 7C) and overall protein content (Figure 7D) in mutant hearts.

**Figure 7.**
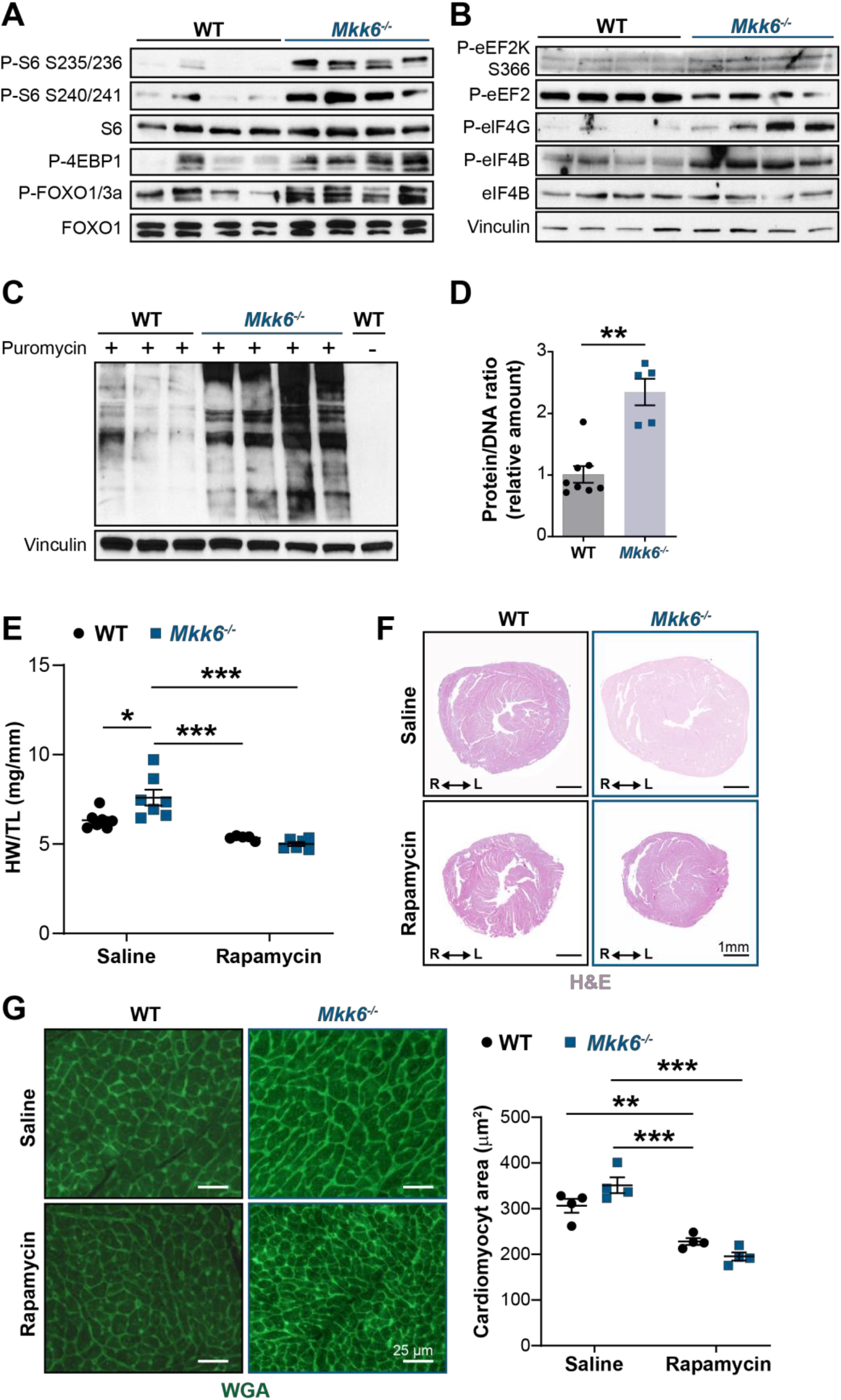
Hyperactivation of mTOR signaling drives cardiac hypertrophy in *Mkk6^-/-^* mice. (**A**, **B**) Immunoblot analysis of mTOR signaling pathway activity (**A**) and activation status of translation factors (**B**) in heart lysates from 9-week-old WT and *Mkk6^-/-^* mice. (**C**) *In vivo* measurement of protein synthesis. Mice were injected intraperitoneally with 0.040 μmol g ^-1^ puromycin dissolved in 100 μl PBS. Exactly 30 min after injection, tissues were extracted and frozen in liquid N_2_ for immunoblot analysis with anti-puromycin antibody. (**D**) Protein content of WT (*n*=8) and *Mkk6^-/-^* (*n*=5) hearts measured as the protein-DNA ratio. Mann-Whitney test. (**E**) Heart weight to tibia length ratio in WT (*n*=5-7) and *Mkk6^-/-^* (*n*=6-7) mice after rapamycin treatment. Mice received daily intraperitoneal injections with rapamycin (2 mg kg ^-1^ per day) or vehicle from weeks 4 to 9 after birth. 2-way ANOVA followed by Tukey’s posttest. (**F**) Representative H&E-stained transverse heart sections after treatment. Scale bars: 1 mm. (**G**) Representative FITC-WGA staining and corresponding quantification of cardiomyocyte cross-sectional area from WT (*n*=4) and *Mkk6^-/-^* (*n*=4) mice hearts after rapamycin treatment. 2-way ANOVA followed by Tukey’s post test. Scale bars: 25 μm. Data in D, E and G are mean ± SEM. \**P*<0.05; \*\**P*<0.01; \*\*\**P*<0.001.

**Figure 8.**
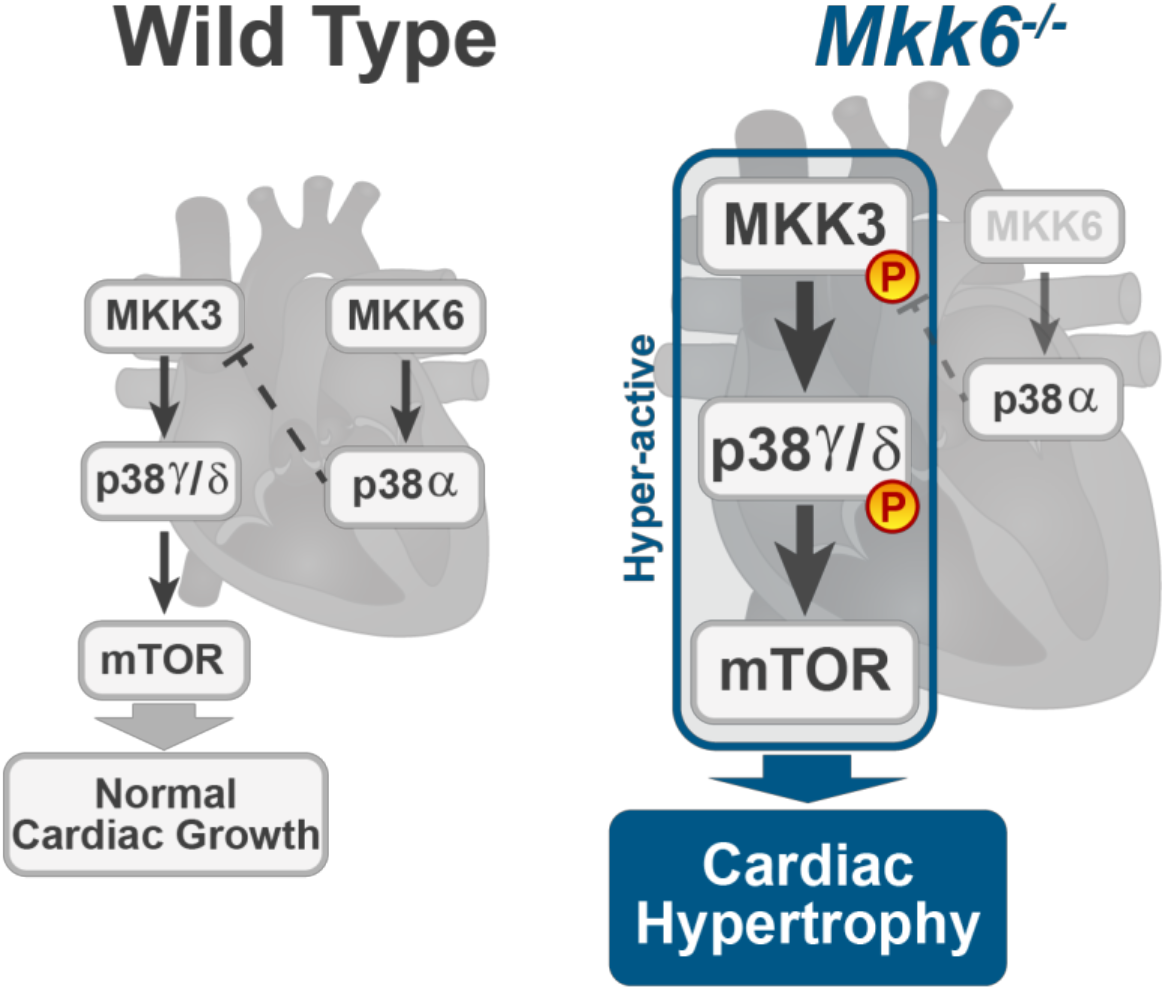
Model for p38γ/δ activation mediated cardiac hypertrophic growth. In a physiological context, MKK3-p38γ/δ pathway promotes normal cardiac growth through the activation of mTOR signaling pathway. MKK6 deficiency stimulates the hyperactivation of MKK3-p38γ/δ and the consequent increase in mTOR activity, which drives increased cardiac hypertrophy.

We next examined the extent to which cardiac hypertrophy in *Mkk6^-/-^* mice is mediated by increased mTOR signaling. We blocked mTOR activation by daily intraperitoneal injection of rapamycin, a potent and specific small molecule mTOR inhibitor, from 3 to 9 weeks of age. This treatment was sufficient to bring the heart size (HW/TL) of *Mkk6^-/-^* animals close to that of WT age-matched controls, which corresponded with a robust reduction in cardiomyocyte cross-sectional area in *Mkk6^-/-^* mice (Figure 4), altogether suggesting that the cardiac hypertrophy in *Mkk6^-/-^* mice results from hyperactivation of mTOR signaling.

## DISCUSION

The present study provides several independent lines of evidence supporting a critical role for the MKK3/6–p38γ/δ signaling pathway in the development of cardiac hypertrophy. The hypertrophy developed seems to be physiological in young animals, with normal or even increased cardiac function at baseline. However, aging induced the development of cardiac disfunction and premature death. Extensive analysis with different knockout mouse models shows that in the absence of MKK6, the MKK3-stimulated p38γ/δ kinases become hyperactivated and induce enhanced postnatal hypertrophic growth through the mTOR pathway. Our results also confirm that the two main up-stream p38 activators are strongly biased toward the activation of specific p38 MAPK isoforms in the heart, with p38γ/δ mainly regulated by MKK3 and p38α by MKK6, at least in homeostatic conditions.

Previous work has implicated multiple p38 isoforms in disease models of pathologic hypertrophy (Nikolic *et al*., 2020; Romero-Becerra *et al*., 2020), whereas p38γ and p38δ isoforms appeared to be involved primarily in regulating postnatal physiological cardiac growth and the metabolic switch during cardiac early postnatal development (Gonzalez-Teran *et al*., 2016). Such reports, however, have not addressed how different p38 MAPK isoforms are regulated and the relative contributions of MKK3 and MKK6 to cardiac hypertrophic phenotypes have so far remained unclear. Indeed, either overexpression of constitutively active or dominant negative forms of MKK3/6 *in vitro* yields cardiac hypertrophy (Braz *et al*, 2003; Streicher *et al*, 2010; Zechner *et al*, 1997). Our *in vivo* data support a model wherein MKK3 activity promotes hypertrophic growth. We show that MKK3 hyperactivation resulting from MKK6 deletion leads to cardiac hypertrophy while MKK3 genetic ablation results in reduced postnatal cardiac growth. We also demonstrate that deficiency of MKK6 in cardiomyocytes is sufficient to result in cardiac hypertrophy, attributable to both, MKK6 direct function in inducing hypertrophy via p38-mediated signaling as well as its role as a negative regulator of MKK3 activity. This negative regulation may be a direct action of MKK6 or may be mediated by p38α, whose phosphorylation levels are strongly decreased in *Mkk6^-/-^* hearts. The last possibility is further supported by evidence indicating that p38α might negatively regulate the MKK3/6 upstream kinase MAP3K TAK1 (Singhirunnusorn *et al*, 2005). Indeed, we have found that in other tissues, lack of p38α induces p38γ and p38δ activation (Matesanz *et al*, 2018). The negative feedback activity of p38α is consistent with MKK3 hyperphosphorylation identified in *Mkk6^-/-^* mice. Our results provide the first demonstration that MKK3 preferentially regulates p38γ and p38δ activation, whereas MKK6 is responsible for p38α regulation.

MKK6-deficient mice show p38γ/δ hyperactivation and impaired p38α activation, possibly implicating any of these isoforms in the observed phenotypes. Through the use of multiple *in vivo* models, we show that hyperphosphorylated MKK3 promotes cardiac hypertrophic growth through activation of p38γ- and p38δ-mediated mTOR signaling in the heart. This is consistent with our earlier work showing an essential role for these p38 isoforms in mTOR-dependent of physiologic and pathological cardiac hypertrophy (Gonzalez-Teran *et al*., 2016), as well as with previous reports indicating that p38α does not mediate hypertrophic responses in animal models of pressure-overload cardiac hypertrophy (Nishida *et al*, 2004). This conclusion is corroborated by the reversion of cardiac hypertrophy in MKK6-deficient mice also deficient for cardiac p38γ or p38δ, and further corroboration comes from the ability of the mTOR inhibitor rapamycin to prevent cardiac hypertrophy during early postnatal cardiac development in MKK6-deficient mice. These findings correlate with impaired p38γ and p38δ activation and reduced postnatal hypertrophic growth in MKK3-deficient mice.

The p38γ- and p38δ-mediated control of cardiac hypertrophy during postnatal cardiac development resides in cardiomyocytes (Gonzalez-Teran *et al*., 2016). Accordingly, lack of MKK6 in skeletal muscle or cardiomyocytes yields the same phenotype as global MKK6 deficiency. Furthermore, the blockade of cardiac hypertrophy in MKK6-deficient mice upon deletion of p38δ specifically in striated muscle indicates that cardiac p38δ lies downstream of MKK6 in the signaling pathway controlling hypertrophic growth. Interestingly, reversion of the MKK6-deficient phenotype upon p38δ deletion was greater than that achieved upon deletion of p38γ (Figure 6), consistent with the reported dominance of p38δ in regulating cardiac hypertrophic growth (Gonzalez-Teran *et al*., 2016).

Young MKK6-deficient mice develop a cardiac hypertrophy that could be classified as physiological, characterized by a proportionate increase in heart size with maintenance of a normal cardiac structure. Moreover, cardiac function was normal, with increased cardiac output and stroke volume, and there was no evidence of fibrosis nor re-expression of the cardiac stress fetal gene program. Physiologic cardiac hypertrophy is an adaptive response that increases ventricular mass while maintaining or enhancing cardiac function (Kang, 2006; Nakamura & Sadoshima, 2018). However, it has increasingly become appreciated that sustained activation of the pathways that drive this beneficial response can ultimately result in pathological remodeling and associated sudden cardiac death (Condorelli *et al*, 2002; Lauschke & Maisch, 2009; Matsui *et al*, 2002; McMullen *et al*., 2004a; Oldfield *et al*., 2020; Shioi *et al*, 2002). In agreement with this, the cardiac hypertrophy observed in *Mkk6^-/-^* mice becomes deleterious with age, compromising the cardiac function and likely contributing to the reduced survival observed in these mice. Our results identify a key role for the MKK3/6-p38γ/δ pathway in the development of cardiac hypertrophy and illustrates how, depending on the stimulus, the activation of the same pathway can promote the progression from a physiological to a pathological phenotype.

The activation of p38α pathway has been linked to several diseases, suggesting that this pathway could represent a target for their treatment. However, the results from the clinical trials have been disappointing so far (Canovas & Nebreda, 2021). p38α inhibitors have been the more profoundly studied. However, these studies usually do not consider the possible undesired effect of p38α inhibition upon the other p38 pathway kinases, which could be among the reasons of failure in the outcomes. Our results show an example of how p38α inhibition leads to an unexpected activation of MKK3-p38γ/δ, having deleterious effects in the heart in the long-term. Our finding suggests that treatment strategies using longstanding p38α inhibition should consider the potential cardiovascular risk among the possible secondary effects of the treatment.

## MATERIALS AND METHODS

### Animal preparation

*Mkk3^−/−^* mice (B6.129-*Map2k3^tm1Flv^*) (Lu *et al*., 1999; Wysk *et al*., 1999) and *Mkk6^-/-^* mice (B6.129-*Map2k6^tm1Flv^*) (Brancho *et al*., 2003) were as previously described. Mice with a germ-line mutation in the *Map2k6* gene and *LoxP* elements inserted into two introns (*Map2k6^LoxP^*) were generated as previously described ^24^. To generate mice lacking MKK6 or p38δ in striated muscle, *Map2k6^LoxP^* or p38δ-negative (B6.129-Mapk13tm1) mice were crossed with the FVB-Tg(Ckmm-cre)5Khn/J line on the C57BL/6J background (Jackson Laboratory). Mice lacking MKK6 in cardiomyocytes were generated by crossing *Map2k6^LoxP^* mice with the Tg(Myh6-cre)2182Mds line on the C57BL/6J background (Jackson Laboratory). The p38γ-negative line (B6.129-Mapk12tm1) was crossed with the *Mkk6^−/−^* line (B6.129-*Map2k6^tm1Flv^*) to generate double knockout mice. Likewise, mice lacking p38δ in striated muscle were crossed with the *Mkk6^−/−^* (B6.129-*Map2k6^tm1Flv^*) line. Genotype was confirmed by PCR analysis of genomic DNA. For signaling studies, animals were killed by cervical dislocation. For rapamycin treatment, mice received daily intraperitoneal injections with rapamycin (LC Laboratories, R-5000) (2 mg kg^-1^ per day) or vehicle (0.25 % polyethylene glycol (Sigma), 0.25 % Tween-20 (Sigma) in PBS); injections started at 4 weeks of age and continued until 9 weeks of age, when heart size was analyzed by echocardiography. All animal procedures conformed to EU Directive 86/609/EEC and Recommendation 2007/526/EC regarding the protection of animals used for experimental and other scientific purposes, enacted under Spanish law 1201/2005.

### Computed tomography scan

Computed tomography (CT) studies were performed with a small-animal PET/CT scanner (nanoScan, Mediso, Hungary). For the acquisition, mice were anesthetized using isoflurane 2% and 1.8 L/min oxygen flow. Ophthalmic gel was placed in the eyes to prevent drying. CT was acquired using an X-ray beam current of 178 μA and a tube voltage of 55 kVp with 360 projections of 500 ms in a helical scan with pitch 1 and binning 1:4. CT image was reconstructed using a Ramlack algorithm with a final resolution of 0.078 mm^3^.

### Histology

Tissue samples were fixed in 10% formalin for 48h, dehydrated, and embedded in paraffin. Sections (8 μm) were cut and stained with hematoxylin and eosin (American Master Tech Scientific). Fibrosis was assessed by Picrosirius red staining (Sigma) and the positive area for fibrosis was quantified with Image J software(Schneider *et al*, 2012). For wheat germ agglutinin (WGA) immunofluorescence, 8 μm heart sections were prepared, washed in PBS, incubated for 2h in WGA-Alexa 488 lectin (Invitrogen, Carlsbad, CA, USA), and washed and mounted in anti-fade reagent. Four images (×20) were taken from each heart, and the areas of 100–200 cross-sectionally oriented cardiomyocytes were measured and analyzed with Image J software(Schneider *et al*., 2012).

### Echocardiography

Mice were anesthetized by inhalation of isoflurane and oxygen (1.25 % and 98.75 %, respectively), and echocardiography was performed with a 30-MHz transthoracic echocardiography probe. Images were obtained with the Vevo 2100 micro-ultrasound imaging system (VisualSonics, Toronto, Canada). Short-axis, long-axis, B-mode and two-dimensional M-mode views were obtained. Scans were conducted by two experienced researchers blinded to the mouse genotype. Measurements of left parasternal long and short axes and M-mode images (left parasternal short axis) were obtained at a heart rate of 500–550 b.p.m. LV end-diastolic diameter (LVEDD), LV end-systolic diameter (LVESD), and wall thickness were measured from M-mode tracings, and the average of three consecutive cardiac cycles is reported. The LV fractional shortening percentage was calculated as ([LVEDD–LVESD]/LVEDD) × 100. MRI of lung was performed with a 7-T Agilent scanner (Agilent, Santa Clara, CA, USA) equipped with a DD2 console and an actively shielded gradient set (205/120 insert of maximum 130 mT m^-1^ gradient strength). To enhance the signal-to-noise ratio during image acquisition, we used a combination of a 72-mm inner diameter quadrature birdcage TX volume coil (Rapid Biomedical GmBH, Germany) and an actively detuning 30-mm flexible customized surface RX coil (Neos Biotec, Pamplona, Spain). After acquisition of a tripilot gradient-echo image, a gradient-echo sequence without gating was used to acquire oblique coronal slices (1-2 slices) and axial slices (7-10 slices covering the entire lung, 72-s acquisition time per slice) using the following parameters: TR/TE=6.7/2.2 ms, flip angle=10 degree, bandwidth=100kHz, field of view=3 × 3 cm, matrix=256 × 128, slice thickness=1 mm (ref. 40). These images were used to determine interventricular septum and left ventricle posterior wall thicknesses and left ventricle corrected mass; the short-axis M-mode quantification was chosen as the most representative. Function was estimated from the ejection fraction and fractional shortening obtained from M-mode views by a blinded echocardiography expert. For ejection fraction measurements, a long- or short-axis view of the heart was selected to obtain an M-mode registration in a line perpendicular to the left ventricular septum and posterior wall at the level of the mitral chordae tendineae.

### Immunoblot analysis

Tissue extracts were prepared in Triton (20 mM Tris (pH 7.4), 1 % Triton X-100, 10 % glycerol, 137 mM NaCl, 2 mM EDTA, 25 mM β-glycerophosphate, 1 mM sodium orthovanadate, 1 mM phenylmethylsulfonyl fluoride, and 10 μg ml^-1^ aprotinin and leupeptin). Extracts (20–50 μg protein) and immunoprecipitates (prepared from 0.5-2 mg) were examined by immunoblot. For immunoprecipitation assays, heart extracts were incubated with 1-4 μg of a specific antibody coupled to protein-G-Sepharose. After incubation overnight at 4 °C with agitation, the captured proteins were centrifuged at 10,000g, the supernatants collected, and the beads washed four times in PBS1X. Beads were boiled for 5 min at 95 °C in 10 μl sample buffer. Extracts and immunoprecipitates were examined by SDS–PAGE and blotted with antibodies to the following targets: p38γ and p38δ(Sabio *et al*, 2005; Sabio *et al*, 2004) at 1 μg ml^-1^; vinculin (Sigma); puromycin (Millipore clone 12D10); phospho-MKK3 (Ser189)/MKK6 (Ser207), MKK3, MKK6, phospho-p38 MAPK (Thr180/Tyr182), phospho-mTOR (Ser2481), mTOR, phospho-p70S6 kinase (Thr 389) (108D2), p70S6 kinase, phospho-S6 (Ser 235/236) (D57.2.2E), phospho-S6 (Ser 240/244) (61H9), S6 ribosomal protein, phospho-FoxO1 (Thr24)/FoxO3a (Thr32), phospho-eEF2 (Thr56), phospho-eIF4G (Ser1108), phospho-eIF4B (Ser422), eIF4B, phospho-4EBP1 (Thr37/46), and 4EBP1, all at a 1:1000 dilution. Immunocomplexes were detected by enhanced chemiluminescence (GE Healthcare Lifesciences).

### *In vivo* protein synthesis assay

For all *in vivo* measurements of protein synthesis, mice were injected intraperitoneally with 0.040 μmol g ^-1^ puromycin dissolved in 100 μl PBS. Exactly 30 min after injection, tissues were extracted and frozen in liquid N_2_ for subsequent immunoblot analysis of protein-incorporated puromycin.

### Blood pressure and heart rate measurements

Blood pressure and heart rate in mice was measured using the noninvasive tail-cuff method(Kubota *et al*, 2006). The measures were performed in conscious mice placed in a BP-2000 Blood Pressure Analysis System (Visitech Systems). 10 preliminary measurements and 10 actual measurements were recorded and the average of the 10 actual measurements used for analysis. The animals were trained for 4 consecutive days prior the actual measurements were registered. All the measurements were taken at the same time of the day.

### RT-qPCR

RNA 500ng – extracted with RNAeasy Plus Mini kit (Qiagen) following manufacturer instructions - was transcribed to cDNA, and RT-qPCR was performed using Fast Sybr Green probe (Applied Biosystems) and the appropriate primers in the 7900 Fast Real Time thermocycler (Applied Biosystems). Relative mRNA expression was normalized to *Gapdh* mRNA measured in each sample. *Fn1* Fw: ATGTGGACCCCTCCTGATAGT, Rev: GCCCAGTGATTTCAGCAAAGG; *Colla1* Fw: GCTCCTCTTAGGGGCCACT, Rev: CCACGTCTCACCATTGGGG; *Col3a1* Fw: CTGTAACATGGAAACTGGGGAAA, Rev: CCATAGCTGAACTGAAAACCACC; *Nppa* Fw: GCTTCCAGGCCATATTGGAG, Rev: GGGGGCATGACCTCATCTT; *Nppb* Fw: GAGGTCACTCCTATCCTCTGG, Rev: GCCATTTCCTCCGACTTTTCTC; *Acta-2* Fw: CCCAAAGCTAACCGGGAGAAG, Rev: CCAGAATCCAACACGATGCC; *Myh7* Fw: ACTGTCAACACTAAGAGGGTCA, Rev: TTGGATGATTTGATCTTCCAGGG; *Gapdh* Fw: TGAAGCAGGCATCTGAGGG, Rev: CGAAGGTGGAAGAGTGGGA

### Statistical analysis

Results are expressed as mean ± SEM. A difference of *P<0.05* was considered significant. Gaussian (normal) distribution was determined using the Shapiro-Wilks normality test. For normally distributed populations, differences between groups were examined for statistical significance by two-tailed Student *t*-test (2 groups) and 1-way ANOVA followed by Tukey post-test (3 or more groups). To test the respective roles of treatment or age and genotype, a 2-way ANOVA was performed. Tukey or Sidak post-test were subsequently employed when appropriate. For data that failed normality testing, Mann-Whitney test (2 groups), or Kruskal-Wallis with Dunn post-test (3 or more groups) was performed. Gehan-Breslow-Wilcoxon test was used to assess significance in the Kaplan–Meier survival analysis.

## Data Availability

All data generated or analysed during this study are included in the manuscript and supporting file; Source Data files have been provided for Figures 1-7

N/A

## Ethics

Human Subjects: No Animal Subjects: Yes Ethics Statement: This study was performed in strict accordance with the recommendations in the Guide for the Care and Use of Laboratory Animals of the National Institutes of Health. All animal procedures conformed to EU Directive 86/609/EEC and Recommendation 2007/526/EC regarding the protection of animals used for experimental and other scientific purposes, enacted under Spanish law 1201/2005. All of the animals were handled according to approved institutional animal care and use committee protocols (PROEX-215/18) of the Comunidad de Madrid.

## ACKNOWLEDGEMENTS

We thank S. Bartlett and F. Chanut for English editing. We are grateful to R.J. Davis, A. Padmanabhan, M. Costa and C. López-Otín for critical reading of the manuscript. We thank A. C. Silva for help with figure editing and design. This work was funded by a CNIC Intramural Project Severo Ochoa (Expediente 12-2016 IGP) to GS and JJ and MINECO-FEDER, PID2019-104399RB-I00 to GS. BGT was a fellow of FPI Severo Ochoa CNIC Program (SVP-2013-067639) and is an American Heart Association Postdoctoral Fellow (18POST34080175). R.R.B is a fellow of the FPU Program (FPU17/03847). The following grants provided additional funding: GS is granted by funds from European Regional Development Fund (ERDF): EFSD/Lilly European Diabetes Research Programme Dr Sabio, Fundación AECC PROYE19047SABI and Comunidad de Madrid IMMUNOTHERCAN-CM B2017/BMD-3733; US National Heart, Lung, and Blood Institute (R01 Grant HL122352), Fondos FEDER, Madrid, Spain, and Fundación Bancaria “La Caixa (project HR19/52160013); Fundación La Marató TV3: Ayudas a la investigación en enfermedades raras 2020 (LA MARATO-2020); and Instituto de Salud Carlos III to JJ. IN. was funded by EFSD/Lilly grants (2017 and 2019), the CNIC IPP FP7 Marie Curie Programme (PCOFUND-2012-600396), EFSD Rising Star award (2019), JDC-2018-Incorporación (MIN/JDC1802). The CNIC is supported by the Instituto de Salud Carlos III (ISCIII), the Ministerio de Ciencia e Innovación (MCIN) and the Pro CNIC Foundation)

## AUTHOR CONTRIBUTIONS

G.S. conceived this project. B.G-T. and R.R-B. equally performed the primary experiments, acquired, prepared figures, and analyzed the data. G.S., B.G-T. and R.R-B. designed, developed the hypothesis and wrote the manuscript with input from all authors. B.G-T., R.R-B. A.M., E.M., L.S., I.N., V.M-R., F.M.C.U., A.M.S., M.E.R., L.L-V., L.J.J-B, J.J., D.F-R. and V.B. help to perform some of the experiments and acquired data and critically revised the manuscript. G.S. led and funded the project.

---

**Figure 1 – video supplement 1. Ataxia and hunched posture in *Mkk6^-/-^* mice.** Video showing the movement of 18-month-old WT (left) and *Mkk6^-/-^* (right) mice. WT: wild type.

**Figure 3 – figure supplement 1.**
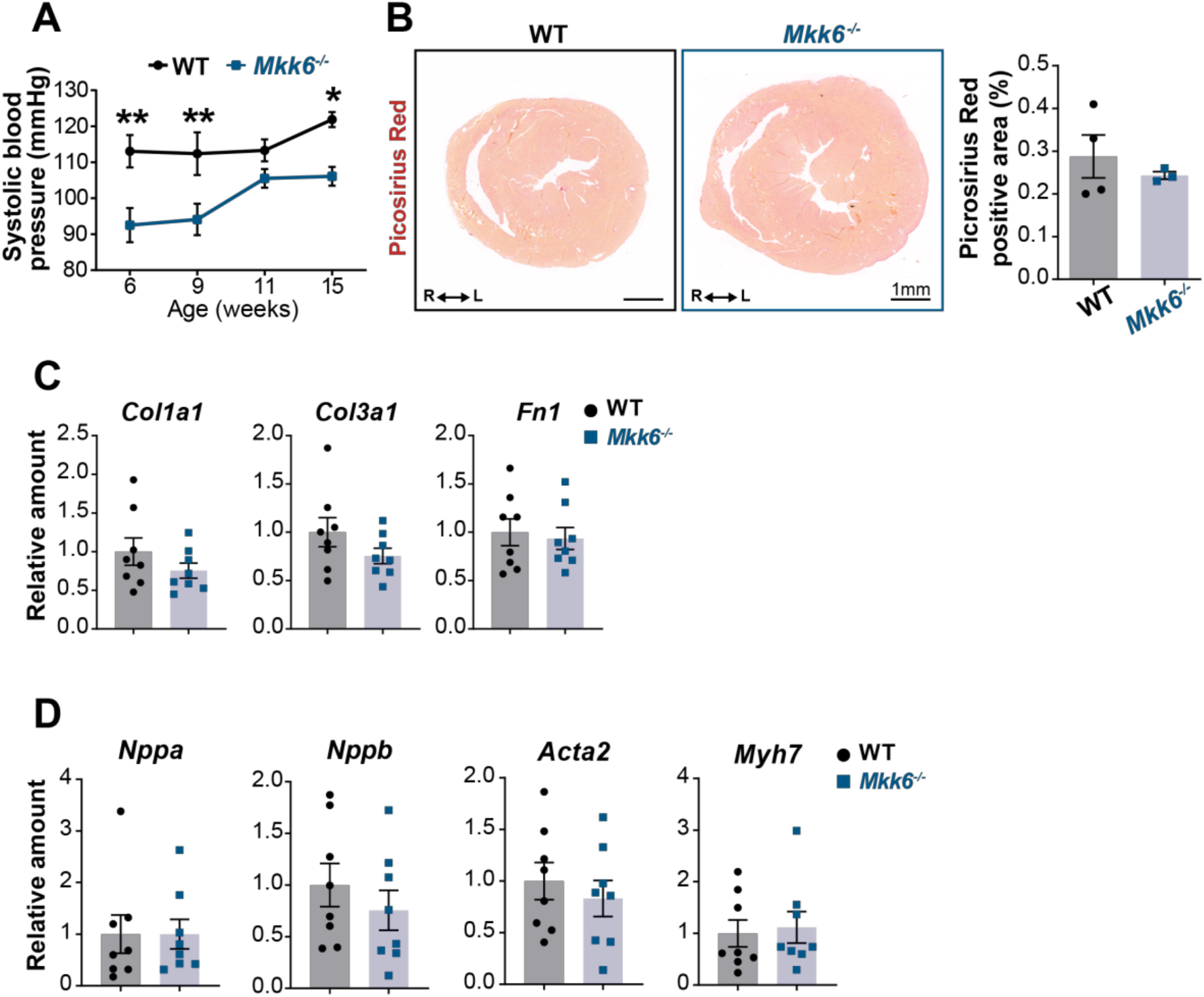
Evaluation of hallmarks of pathological cardiac hypertrophy in young *Mkk6^-/-^* mice. (**A**) Systolic blood pressure in WT (*n*=5) and *Mkk6^-/-^* (*n*=4-7) mice at the indicated times after birth. 2-way ANOVA followed by Sidak’s post test. (B) Picrosirius red staining and quantification of cardiac fibrosis in 9-week-old WT (*n*=4) and *Mkk6^-/-^* (*n*=3) mice. Unpaired *t*-test. Scale bars: 1mm. (**C-D**) Cardiac gene expression of fibrosis (C) and cardiac stress (D) markers in 9-week-old WT (*n*=8) and *Mkk6^-/-^* (*n*=8). Unpaired *t*-test or Mann-Whitney test. Data in A-D are mean±SEM. *P<0.05. **P<0.001

**Figure 4 - figure supplement 1.**
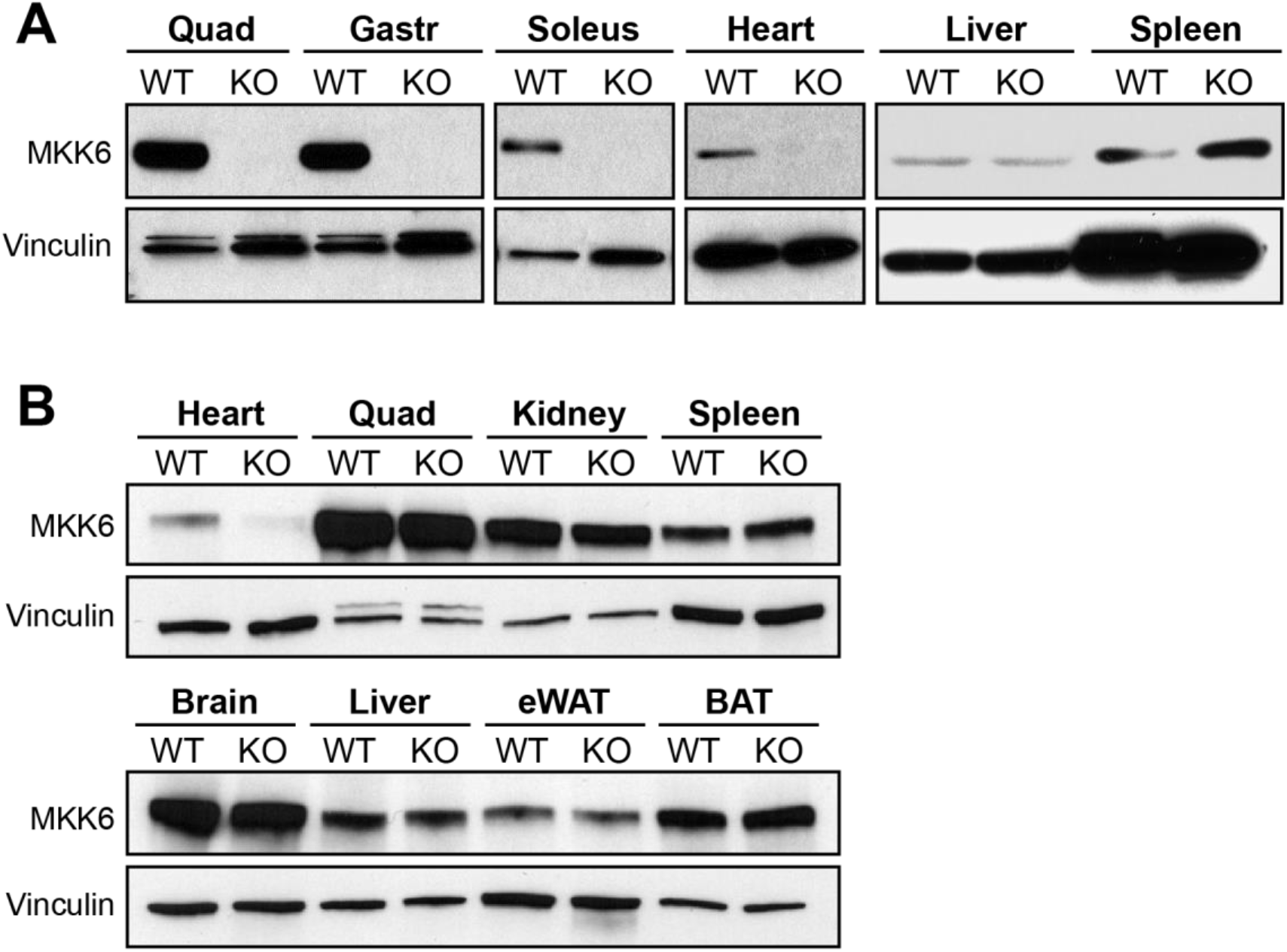
Tissue-specific MKK6 deletion in Mkk6^MCK-KO^ and Mkk6^αMHC-KO^ mice. (**A**) Confirmation of MKK6 deletion in tissue lysates from Mkk6^MCK-KO^ (KO) mice in quadriceps (Quad), gastronemius (Gastr), soleus and heart, but not in spleen and liver, and not in MCK-Cre (WT) tissues. (**B**) Confirmation of specific MKK6 deletion in the heart of Mkk6^αMHC-KO^, but not in lysates of αMHC-Cre (WT) and Mkk6^αMHC-KO^, (KO) quadriceps (Quad), kidney, spleen, brain, liver, epididymal white adipose tissue (eWAT) and brown adipose tissue (BAT).

